# CLEC18A interacts with sulfated GAGs and controls clear cell renal cell carcinoma progression

**DOI:** 10.1101/2024.07.08.602586

**Authors:** Gustav Jonsson, Maura Hofmann, Stefan Mereiter, Lauren Hartley-Tassell, Irma Sakic, Tiago Oliveira, David Hoffmann, Maria Novatchkova, Alexander Schleiffer, Josef M. Penninger

**Affiliations:** Institute of Molecular Biotechnology of the Austrian Academy of Sciences, IMBA, Dr. Bohr-Gasse 3, 1030 Vienna, Austria; Vienna BioCenter PhD Program, Doctoral School of the University of Vienna and Medical University of Vienna, 1030, Vienna, Austria; Eric Kandel Institute, Department of Laboratory Medicine, Medical University of Vienna, Vienna, Austria; Institute for Glycomics, Griffith University, Southport, QLD, Australia; Department of Medical Genetics, Life Sciences Institute, University of British Columbia, Canada; Helmholtz Centre for Infection Research, Braunschweig, Germany

**Keywords:** C-type lectin, glycosaminoglycan, proteoglycan, proximal tubule, clear cell renal cell carcinoma

## Abstract

C-type lectins are a large family of proteins with essential functions in both health and disease. In cancer, some C-type lectins have been found to both promote and inhibit tumor growth, but many of the C-type lectins still remain uncharacterised in a tumor context. Therefore, there is growing interst in further elucidating the mechanisms with which C-type lectins control tumor growth. Here, we report a key role of the CLEC18 family of C-type lectins in the progression of clear cell renal cell carcinoma (ccRCC). The CLEC18 family is conserved across the entire Chordata phylum with recent gene duplication events in humans. We found that CLEC18A is exclusively expressed in the proximal tubule of the kidney and the medial habenula of the brain. We further identified sulfated glycosaminoglycans (GAGs) of proteoglycans as the main CLEC18A ligand, making them unique among C-type lectins. In ccRCC patients, high expression of the *CLEC18* family lectins in the tumor are associated with improved survival. In mouse models of ccRCC, deletion of the mouse ortholog *Clec18a* resulted in enhanced tumor growth. Our results establishes CLEC18A as a novel and critical regulators of ccRCC tumor growth and highlights the potential benefit of modulating *CLEC18* expression in the renal tumor microenvironment.

## Introduction

C-type lectins are a vast superfamily of proteins found across the entire vertebrate lineage with important functions in physiology and disease. The C-type lectins are defined by containing at least one C-type lectin domain (CTLD), functioning as their carbohydrate recognition domain (CRD). Classically, C-type lectins have been studied in the context of both innate and adaptive immunity towards microbial pathogens, but C-type lectins also play key roles in autoimmunity and cancer [1–3]. Overall, C-type lectins show varied evolutionary conservation across vertebrates. Some C-type lectins are specific to human, and others only have weak orthology to commonly used model species making them difficult to study in a disease context [4–6].

However, in the last decades, key insights have been generated into how C-type lectins are involved in cancer. Some C-type lectins, such as Dectin 1, Mincle, MGL and DC-SIGN, have all been found to suppress immunity towards tumor cells through their expression in the myeloid compartment [7–9]. On the other hand, C-type lectins have also been found to be involved in eradicating tumor cells. For example, the C-type lectin NKG2D recognises polymorphic MHC class I related stress induced ligands and serves as a key activating receptor for NK cells [10]. Further, myeloid restricted C-type lectins have been shown to be involved in the clearance of liver metastasis [11] and suppression of hepatocarcinogenesis [12].

Among C-type lectins the secreted CLEC18 family remains poorly characterised. The human *CLEC18* locus, consisting of the paralogs *CLEC18A, CLEC18B* and *CLEC18C*, is located on chromosome 16q22 and encodes for a family of secreted proteins [13]. In viral infections, *CLEC18* has been shown to be downregulated in Hepatitis B infections while upregulated during Hepatitis C infections [14, 15]. Furthermore, CLEC18 associates with TLR3 and thereby stimulates the production of type I and type III interferons following H5N1 influenza A virus exposure [16] and transgenic expression of CLEC18A in *Aedes aegypti* mosquitos significantly reduced dengue virus infectivity [17]. Other than these reports, very little is known about the role of CLEC18 in normal physiology and other diseases, such as cancer. Moreover, the exact carbohydrate ligands for CLEC18 family members remains elusive.

In the present study, we provide a deep phylogenetic analysis of the *CLEC18* family and its orthologs throughout the Chordata phylum. Expression of CLEC18A was mapped to the medial habenula of the brain and the proximal tubule of the kidney. We also resolved the first high-confidence CLEC18A ligands, sulfated glycosaminoglycans (GAGs) on a collection of proteoglycans. Interestingly, *CLEC18* expression was associated with favorable cancer prognosis in ccRCC. Lastly, loss of function studies in mice showed that deletion of *Clec18a* in renal cancer cells resulted in a significant increase in the tumor burden of mice.

## Results

### CLEC18 is a family of conserved gene paralogs restricted to the kidney and brain

To explore evolutionary conservation, we identified CLEC18 orthologs across vertebrates. We found that that human CLEC18A has homologs and paralogs across the Chordata phylum with the earliest identifiable ancestor being lampreys. Unlike some other C-type lectins which lack direct orthology between species [4], CLEC18A instead has different number of paralogs in several species. Lower vertebrates and most mammals only have one copy of CLEC18, whereas some higher species have two or three paralogs, e.g. humans (CLEC18A, CLEC18B, and CLEC18C) (**Supplementary Fig. 1A**), indicating recent gene duplication events. Furthermore, the three human CLEC18 paralogs are almost identical (**Fig. 1A**, **Supplementary Fig. 1B)** and across the entire Chordata phylum the CLEC18 proteins are highly conserved and ordered (**Fig. 1B-D**), when present. For example, no CLEC18 homologs are found in the common model organisms zebrafish and xenopus.

**Figure 1.**
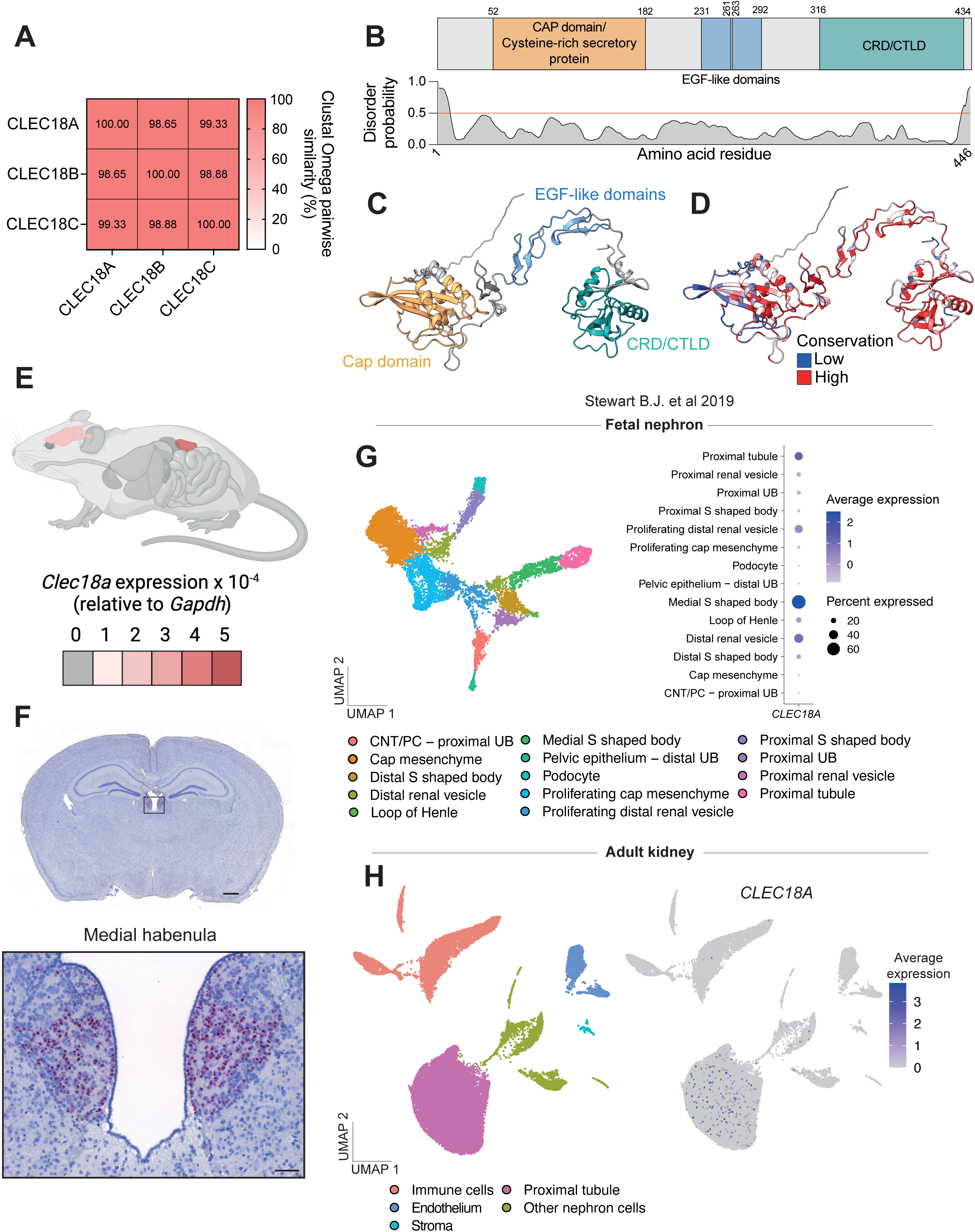
The CLEC18 family are highly conserved proteins and gene expression is restricted to the brain and kidneys. **A.** Similarity matrix between the amio acid sequences of human CLEC18A, CLEC18B and CLEC18C calculated through Clustal Omega pairwise similarity. **B.** Human CLEC18A protein domains and disorder. Disorder is calculated using PrDOS. **C.** AlphaFold2 prediction of human CLEC18A structure with protein domains highlighted. **D.** AlphaFold2 prediction of human CLEC18A structure with overlayed conservation scored calculated by the predicted local distance difference test (pLDDT). **E.** *Clec18a* expression in all organs of mice measured with RT-qPCR. Expression data is shown as an organ heatmap overlayed on a mouse. Expression was predominantely detected in the kidney and brain. Schematic created using BioRender.com. **F.** RNAscope *in situ* hybridization of *Clec18a* on coronal sections of mouse brains. Positive cells display a red color in the insert of the medial habenula. Scale bar for the full coronal brain section = 1 mm. Scale bar for the insert = 100 μm. **G.** Re-analysis of scRNA-seq data from the kidney cell atlas [18] showing clustering of fetal nephron cells and detection of *CLEC18A* in the medial S shaped body and proximal tubules of fetal nephrons as a bubble plot. Abbreviations: UMAP = Unifold Manifold Approximation and Projection, UB = Ureteric bud, CNT = Connecting tubule, PC = Principal cell. **H.** Re-analysis of scRNA-seq data from the kidney cell atlas [18] showing detection of *CLEC18A* in the proximal tubule of adult kidneys. Abbreviations: UMAP = Unifold Manifold Approximation and Projection.

Next, we determined the expression pattern of *Clec18a* mRNA. RT-qPCR of mouse organ revealed highly restricted expression to the brain and kidney, with the kidney exhibiting the highest expression (**Fig. 1E**, **Supplementary Fig. 1C**). RNA fluorescence *in situ* hybridization of the brain revealed that *Clec18a* is almost exclusively expressed in the medial habenula (**Fig. 1F**). Re-analysis of the kidney cell atlas [18] of fetal kidneys revealed that human *CLEC18A* is predominantly expressed in various structures of the fetal nephron (**Supplementary Fig. 2A**). A more detailed assessment of human fetal nephrons revealed that during development *CLEC18A* is most commonly found in the distal renal vesicle, proximal tubules and S-shaped bodies (**Fig. 1G**), the latter of which gives rise to the proximal and distal tubules, and the loop of Henle. In the adult kidney, *CLEC18A* was almost exclusively found in the proximal tubule (**Fig. 1H**, full clustering shown in **Supplementary Fig. 2B**). These data show that CLEC18A is a highly conserved and ordered C-type lectin protein across the Chordata phylum, predominantly found in the medial habenula and proximal tubule of the kidney.

### CLEC18A interacts with glycosaminoglycans (GAGs) on proteoglycans

We next sought to elucidate the ligand of CLEC18 proteins. Phylogenetic analysis and principal component analysis on a multiple sequence alignment of the CRDs of canonical C-type lectins (**Fig. 2A, Supplementary Fig. 3A**) revealed that the CLEC18 paralog family forms its own cluster (**Fig. 2B**), indicating that they have unique CRD sequences and that they might interact with unique ligands. A previous study reported glycoarray analysis with a CLEC18A construct using an array with 611 common N-linked and O-linked glycostructures, but failed to elucidate significant interaction partners [13]. However, despite not reaching significance, the strongest hit was GlcAβ1-6Galβ (**Supplementary Fig. 3B**). Glucuronic acid (GlcA) is primarily used as a building block in glycosaminoglycans (GAGs) and appears in all glycosaminoglycans of proteoglycans except for keratan sulfate. Furthermore, GlcA linked to Galactose (Gal) appears at the start of heparan sulfate and chondroitin sulfate, albeit not with a β1-6 linkage [19]. Assuming that the correct ligand was not included in the original array, these results suggested to us that a proteoglycan GAG could be the ligand.

**Figure 2.**
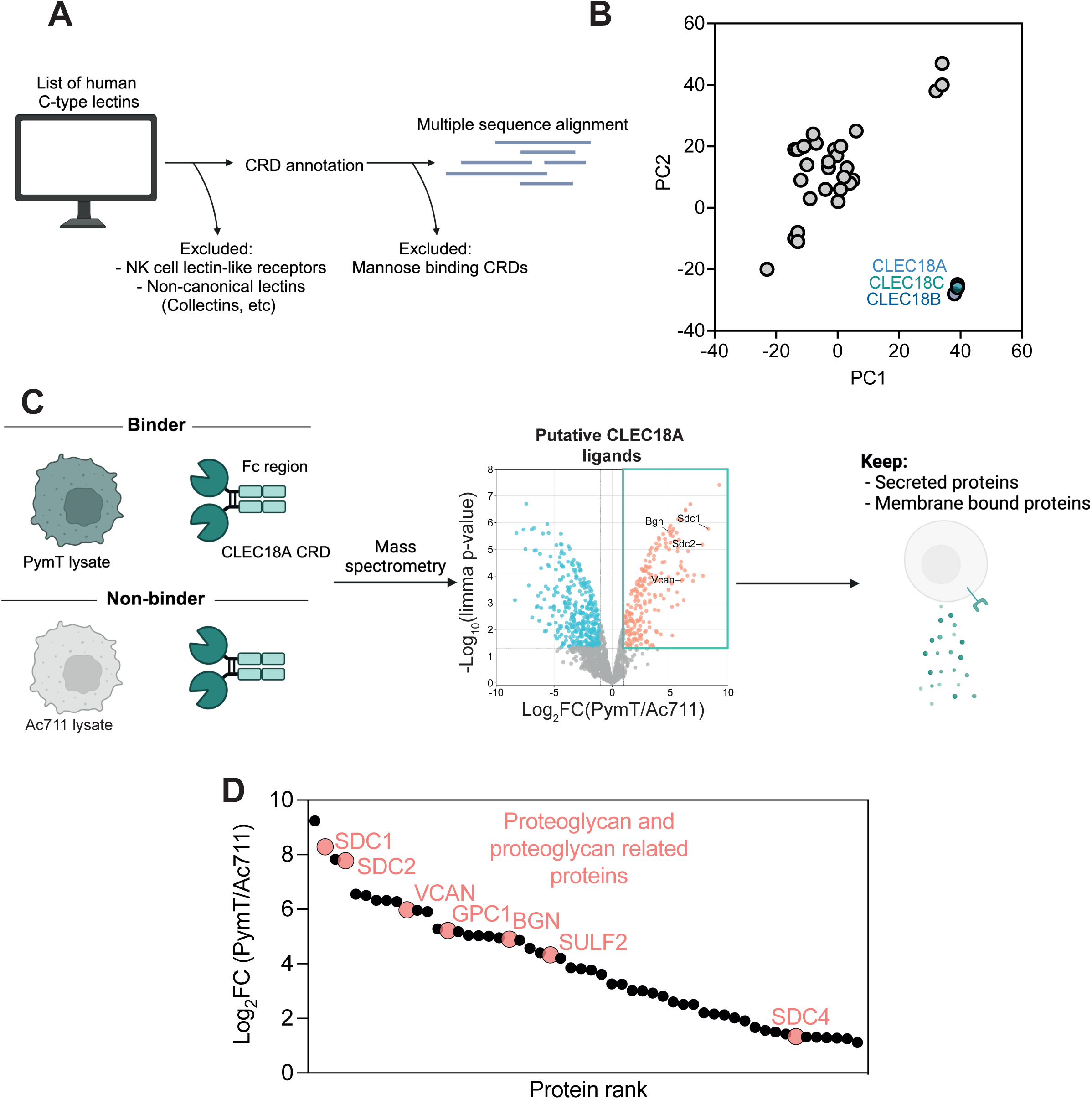
CLEC18A interacts with proteoglycans. **A.** Schematic of CRD annotation of all human C-type lectins and multiple sequence alignment of those sequences. Abbreviations: CRD = Carbohydrate recognition domain. Schematic created using BioRender.com. **B.** Principal component analysis using JalView on the generated multiple sequence alignment of the human CRD sequences. Abbreviations: PC = Principal component **C.** Schematic of CLEC18A-Fc fusion pulldown strategy against two cell lines, one with high binding affinity for CLEC18A (PymT) and one with no binding affinity to CLEC18A (Ac711). Mass spectrometry was performed on pulldowns and putative CLEC18A ligands were filtered for proteins which are either secreted or bound to the membrane. Volcano plot shows upregulated (red) and downregulated (blue) proteins in PymT lysates (binders) vs Ac711 lysates (non-binders). Some proteoglycans are highlighted as interaction partners. Abbreviations: CRD = Carbohydrate recognition domain, Bgn = Biglycan, Sdc1 = Syndecan 1, Sdc2 = Syndecan 2, Vcan = Vesican. P values were calculated with limma-moderated Benjamini–Hochberg-corrected two-sided t-test. Abbreviations: FC = Fold change. Schematic created using BioRender.com. **D.** Filtered secreted and membrane bound ligands hits from **C** shown as a waterfall plot. Proteoglycan and proteoglycan related proteins are highlighted. Abbreviations: FC = Fold change.

To determine if proteoglycans indeed are the ligands of CLEC18, we expressed a fusion between the murine CRD of CLEC18A and an antibody Fc region (CLEC18A-Fc) (**Supplementary Fig. 4A**). The binding of CLEC18A-Fc was then screened against a large panel of cancer cell lines. The strongest binder (PymT, a murine mammary cancer cell line) and one non-binder (Ac711, also a murine mammary cancer cell line) were chosen for downstream analysis (**Supplementary Fig. 4A-B**). Using these two cell lines, multiple putative CLEC18A ligands were identified in a lectin-pulldown approach (**Fig. 2C, Supplementary Table 1**), many of which were Proteoglycans (e.g. Biglycan (*Bgn*), Vestican (*Vcan*), Syndecan 1 (*Sdc1*) and Syndecan 2 (*Sdc2*)). All putative ligands were filtered for proteins that are either secreted or membrane bound since CLEC18A is predicted to be localized to the extracellular matrix, hypothesizing that the exterior of the cell or extracellular space would be the site of interaction. Amongst this annotated list of ligand candidates, multiple proteoglycans and proteoglycan related proteins were found (**Fig. 2D**).

To investigate whether or not the identified interactions in the pulldown actually are GAGs on proteoglycans, or interactions with the protein backbone, we performed an AlphaFold2 Multimer [20–22] screen with human CLEC18A, CLEC18B and CLEC18C against the amino acid sequences of the secreted and membrane-bound hits identified in the pulldown. In this approach we did not identify any highly scoring protein-protein interactions (PPIs), indicating that the interactions are between the CRD of CLEC18 and the GAG of the proteoglycan (**Fig. 3A**). Furthermore, since CLEC18A is expressed in the proximal tubule, we also performed an AlphaFold Multimer screen with CLEC18A, CLEC18B and CLEC18C, but this time against a kidney specific library of proteins consisting of 379 unique amino acid sequences. In this approach, there were also no identified high-confidence PPIs (**Fig. 3B**). To confirm the GAG specificity of CLEC18A, we used our CLEC18A-Fc fusion protein to probe a glycoarray with GAGs and other glyco-structures (**Fig. 3C**). The glycoarray contained a total of 254 glyco-spots. After removing unspecific binding of the Fc-control, 242 structures remained and out of those 29 glycostructures were identified as glyco-ligands of CLEC18A (**Fig. 3D, Supplementary Table 2**). CLEC18A bound to a variety of heparin, heparan sulfate, and chondroitin sulfate (both low and high molecular weight) GAGs. CLEC18A did not interact with any hyaluronan and hyaluronan fragments which are un-sulfated, indicating the need for disaccharide sulfation for CLEC18A docking (**Fig. 3E**). Of note, CLEC14A is the only other C-type lectin which has been reported to interact with proteoglycan GAG chains [23]. Taken together, these data show that CLEC18A interacts with sulfated GAGs on proteoglycans.

**Figure 3.**
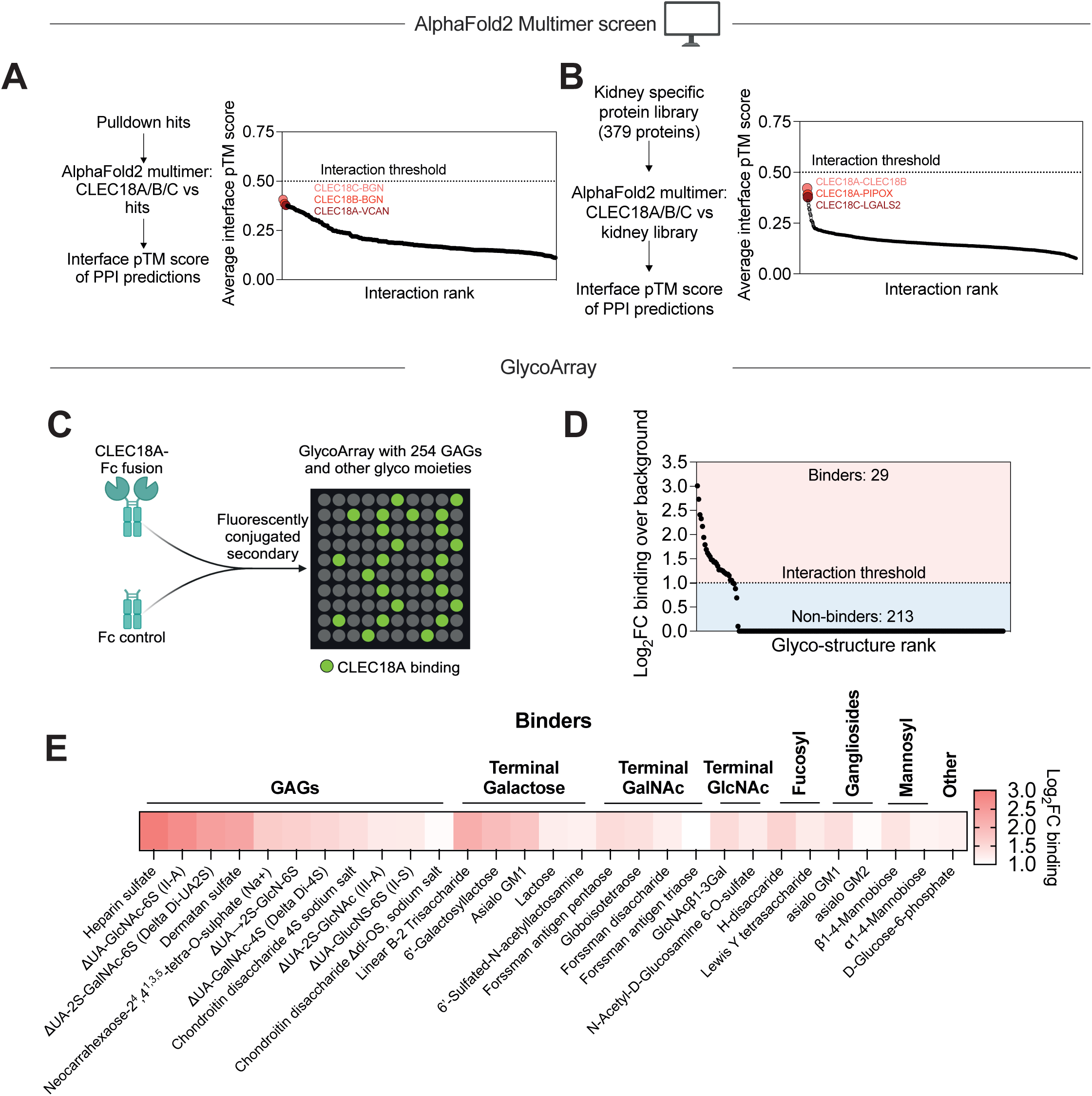
CLEC18A interacts with glycosaminoglycans, and not amino acid residues, on proteoglycans. **A.** AlphaFold2 Multimer screen of CLEC18A, CLEC18B and CLEC18C against the top ligand candidates in the CLEC18A-Fc fusion pulldown. The interaction threshold of interface pTM > 0.5 is not reached for any of the top hits indicating that these are not protein-protein interactions. The three highest scoring predictions are highlighted. Abbreviations: pTM = predicted template modelling, PPI = Protein-protein interaction. **B.** AlphaFold2 Multimer screen of CLEC18A, CLEC18B and CLEC18C against a library of 379 kidney and clear cell renal cell carcinoma specific proteins. The three highest scoring predictions are highlighted. Abbreviations: pTM = predicted template modelling, PPI = Protein-protein interaction. **C.** Schematic for detection of glyco-ligands for CLEC18A using a GlycoArray with 260 GAGs and other glycol moieties. Full list of glyco structures included in the array are available in Supplementary Table 1. Abbreviations: GAG = Glycosaminoglycan. **D.** Waterfall plot of potential glyco-ligands for CLEC18A from the GlycoArray. Structures which displayed binding to the Fc control have been removed. Abbreviations: FC = Fold change. **E.** Heatmap of the 29 binders which were shown to interact with CLEC18A. Abbreviations: GAG = Glycosaminoglycan, GalNAc = N-Acetylgalactosamine, GlcNAc = N-Acetylglucosamine, FC = Fold change.

### Association of CLEC18 expression with kidney and brain cancer patient survival

CLEC18A has been found to be regulated by, or involved in, some viral infections, such as Hepatitis C and Dengue fever [15–17]. Whereas many C-type lectins play a role in host-pathogen interactions, C-type lectins are also commonly involved in cancer [24, 25]. To assess whether *CLEC18A*, *CLEC18B* or *CLEC18C* might have a role in cancer, we first analyzed patient survival data for all cancer types in The Cancer Genome Atlas (TCGA). Interestingly, we found that expression of all three human *CLEC18A*, *CLEC18B* and *CLEC18C* paralogs is correlated with the survival of patients with clear cell renal carcinoma (ccRCC, TCGA abbreviation: KIRC) (**Fig. 4A**). Of note, out of the three kidney cancer types present in the TCGA; KIRC, papillary renal cell carcinoma (KIRP) and chromophobe renal cell carcinoma (KICH), the expression of *CLEC18* gene cluster only correlated with the survival of KIRC patients **(Fig. 4A).** The individual survival curves for these observations are shown in **Fig. 4B-D** for *CLEC18A* and **Supplementary Fig. 5A-F** for *CLEC18B* and *CLEC18C*. Moreover, mRNA expression of the three CLEC18 family members was upregulated in the tumors of KIRC patients but downregulated in KIRP and KICH **(Fig. 4E**). Lastly, *CLEC18* expression was inversely correlated with the survival of patients with Low grade gliomas (LGG) (**Fig. 4A**). These data indicate that, in line with the very restricted expression profiles of the *CLEC18* family members in the brain and kidney, *CLEC18* expression levels are associated with the survival of patients with low grade glioma and clear cell renal cell carcinomas. These data also show that, for KIRC/ccRCC specifically, that the *CLEC18* gene family is upregulated in the tumors and higher expression correlates with better survival for the patients.

**Figure 4.**
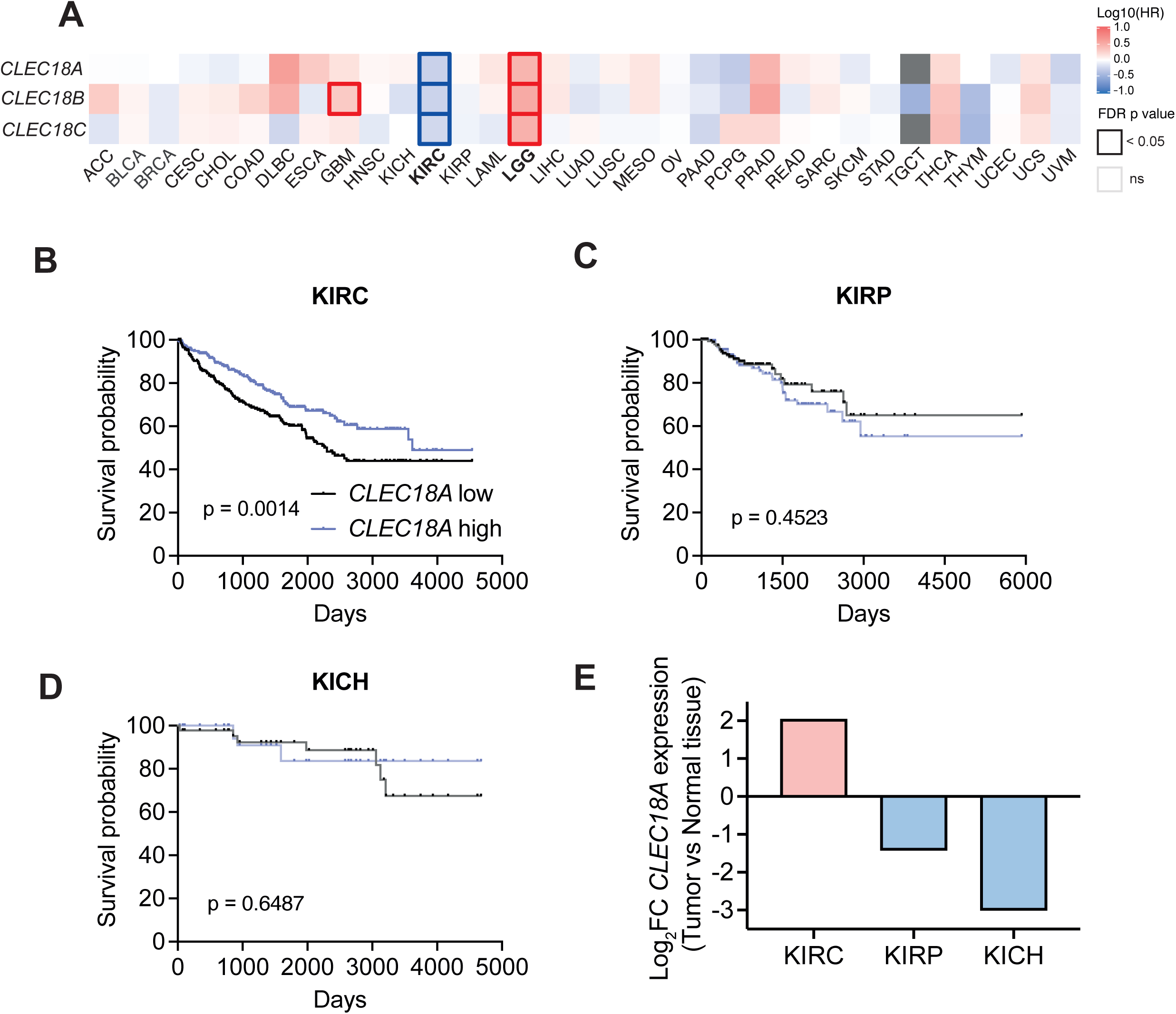
CLEC18A expression correlates with survival in clear cell renal cell carcinoma. **A.** Survival heatmap for high vs low expression (high and low expression attributed based on the median) of *CLEC18A, CLEC18B* and *CLEC18C* for all cancer types in the The Cancer Genome Atlas shown as the hazard ratio. A think outline indicates significant hazard ratios. Cancer types where all three CLEC18 paralogs contribute to a significant shift in survival based on high and low expression are shown in bold; Clear cell renal cell carcinoma (KIRC) and Low grade glioma (LGG). Abbreviations: HR = Hazard ratio, FDR = False discovery rate. **B.** Kaplan meier survival curve for high vs low *CLEC18A* expression (high and low expression attributed based on the median) in KIRC. P value calculated with a Log rank test. Abbreviations: KIRC = Clear cell renal cell carcinoma. **C.** Kaplan meier survival curve for high vs low *CLEC18A* expression (high and low expression attributed based on the median) in KIRP. P value calculated with a Log rank test. Abbreviations: KIRP = Papillary renal cell carcinoma. **D.** Kaplan meier survival curve for high vs low *CLEC18A* expression (high and low expression attributed based on the median) in KICH. P value calculated with a Log rank test. Abbreviations: KICH = Chromophobe renal cell carcinoma. **E.** Expression comparison of *CLEC18A* between tumor and normal tissue for the three kidney cancer types that are present in The Cancer Genome Atlas. Abbreviations: KIRC = Clear cell renal cell carcinoma, KIRP = Papillary renal cell carcinoma, KICH = Chromophobe renal cell carcinoma, FC = Fold change.

### Loss of CLEC18A accelerates kidney cancer cell growth in mice

To further test a potential role of *CLEC18A* in ccRCC, we utilised murine cell lines which only have one CLEC18 paralog, CLEC18A. A murine ccRCC cell line called RAG, which exhibited a high baseline expression of *Clec18a*, was used to generate *Clec18a* knockouts (*Clec18a^-/-^*) using CRISPR-Cas9 (**Fig. 5A**). As a control, we overexpressed *Clec18a* (*Clec18a*^OE^) in an unrelated murine mammary cancer cell line, E0771, which did not have any baseline expression of *Clec18a* (**Fig. 5B**). Deletion or introduction of *Clec18a* had no impact on the *in vitro* growth pattern of either cell line (**Fig. 5C-D**).

**Figure 5.**
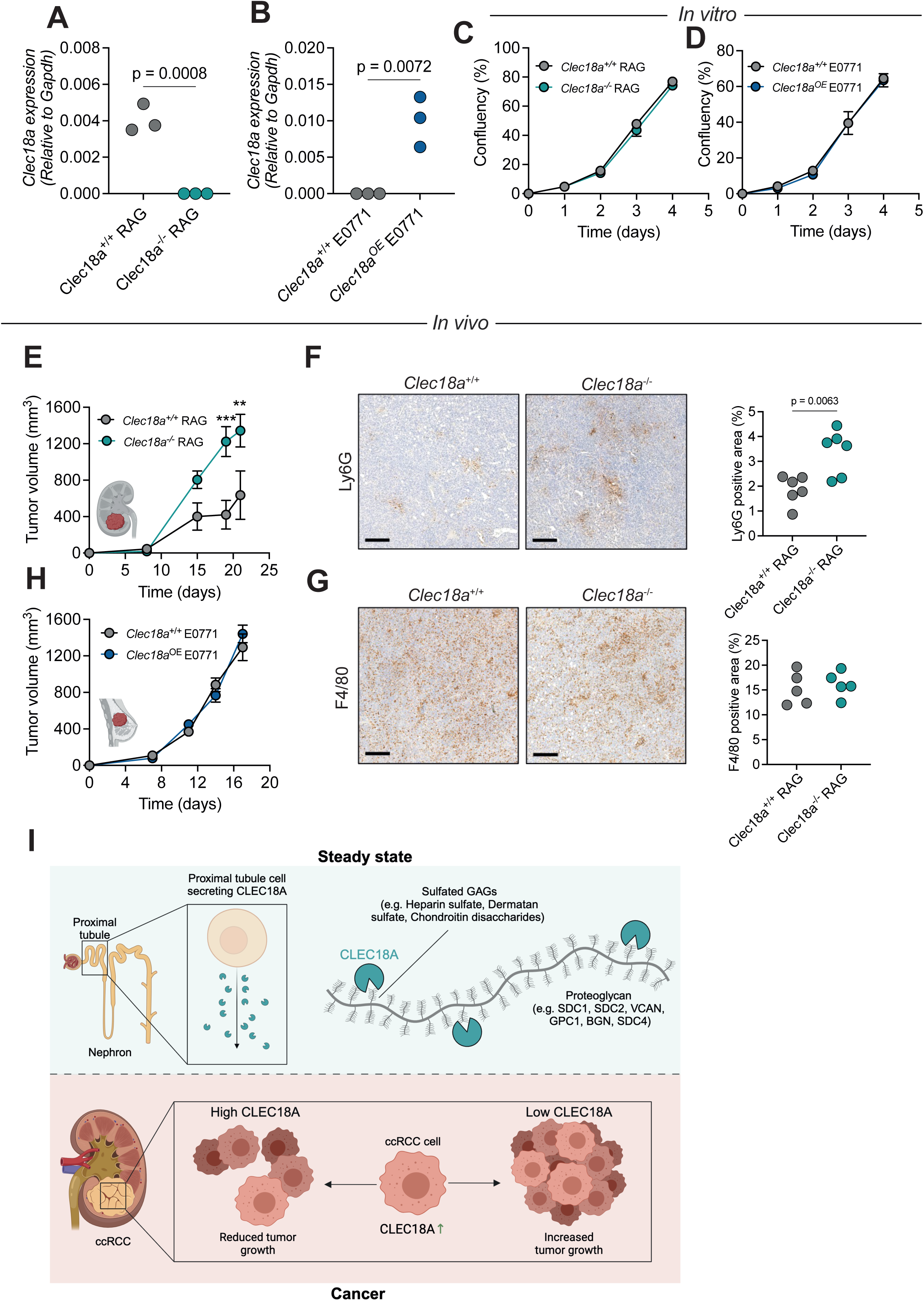
CLEC18A expression restricts tumor progression in murine models of ccRCC. **A.** Validation of *Clec18a* knockout in *Clec18a*^-/-^ RAG cells. P value calculated with a two-sided Student’s t-test. **B.** Validation of *Clec18a* overexpression in *Clec18a*^OE^ E0771 cells. P value calculated with a two-sided Student’s t-test. Abbreviations: OE = overexpression. **C.** *In vitro* growth curves of *Clec18a*^+/+^ and *Clec18a*^-/-^ RAG cells. P values calculated assessed with two-way ANOVA with Šídák’s multiple comparisons test. **D.** *In vitro* growth curves of *Clec18a*^+/+^ (empty vector control) and *Clec18a*^OE^ E0771 cells. P values calculated assessed with two-way ANOVA with Šídák’s multiple comparisons test. **E.** Tumor growth curve kinetics of *Clec18a*^+/+^ and *Clec18a*^-/-^ clear cell renal cell carcinoma cell lines (RAG) in *Rag2*^-/-^ *Il2rg*^-/-^ mice. P values calculated assessed with two-way ANOVA with Šídák’s multiple comparisons test. **F.** Assessment of neutrophil infiltration through Ly6G staining in *Clec18a*^+/+^ and *Clec18a*^-/-^ tumors subcutaneously injected into *Rag2*^-/-^ *Il2rg*^-/-^ mice. Scale bars = 100 μm. P value calculated with a two-sided Student’s t-test. **G.** Assessment of macrophage infiltration through F4/80 staining in *Clec18a*^+/+^ and *Clec18a*^-/-^ tumors subcutaneously injected into *Rag2*^-/-^ *Il2rg*^-/-^ mice. Scale bars = 100 μm. P value calculated with a two-sided Student’s t-test. **H.** Tumor growth curve kinetics of *Clec18a*^+/+^ (empty vector control) and *Clec18a*^OE^ breast cancer cell lines (E0771) in *Rag2*^-/-^ *Il2rg*^-/-^ mice. P values calculated assessed with two-way ANOVA with Šídák’s multiple comparisons test. Abbreviations: OE = Overexpression. **I.** Graphical abstract of the study.

Based on the TCGA data where we found that high *CLEC18A* expression favors survival, we hypothesized that deletion of *Clec18a* in the ccRCC cell line may promote tumor growth. Indeed, genetic inactivation of *Clec18a* promoted tumor growth of RAG ccRCC cancer cells following subcutaneous injection into mice (**Fig. 5E**). Of note, these experiments were performed in *Rag2*^-/-^ *Il2rg*^-/-^ mice which lack T, B and NK cells since the RAG cell line did not grow in immunocompetent mice. Thus, these data indicate that the murine Clec18a hinders tumor progression in a lymphocyte-independent manner. We did find a significantly increased tumor infiltration of Ly6G^+^ neutrophils in *Clec18a*^-/-^ ccRCC tumors (**Fig. 5F**) but no apparent difference in infiltrating F4/80^+^ macrophages (**Fig. 5G**). Overexpression of *Clec18a* had no impact on the tumor progression in the murine breast cancer model in *Rag2*^-/-^ *Il2rg*^-/-^ mice (**Fig. 5H**). In conclusion, our study has shown that the CLEC18 family of proteins are highly conserved. Furthermore, *CLEC18A* is predominantely expressed in the proximal tubule of the kidney and interacts with sulfated GAGs on proteoglycans. Lastly, *CLEC18* is upregulated in the tumor microenvironment of ccRCC and the expression level is positively correlated with survival in both mice and humans (**Fig. 5I**)

## Discussion

C-type lectins are known to play essential roles in cancer ranging from recognition of damaged cells and antigen uptake [26, 27], to elimination of cancer cells through MHC I associated stress signals [10]. In some cases, C-type lectins can also promote tumor growth [9]. The interest for pharmacologically targeting C-type lectins in the clinic for cancer treatments is steadily increasing, but remains a challenging goal due to their complex and numerous mechanisms in malignancies [28]. Therefore, there is an increased need to further our understanding of C-type lectin biology in health and cancer.

In this study we identify a previously poorly characterised C-type lectin family, the CLEC18 family, as novel regulators of ccRCC progression. The CLEC18 gene family has varying numbers of paralogs across the Chordata phylum and higher numbers of paralogs in evolutionary proximity to humans, indicating recent gene duplications and/or other mutational events. Although quite rare across the entire genome, other C-type lectins also display recent evolutionary events. Furthermore, some C-type lectins lack clear orthologs between species and, just like CLEC18A, show species-specific duplications. Prominent examples of copy number differences are the DC-SIGN/SIGN [5, 29] and collectin [6] proteins. Interestingly, for both DC-SIGN/SIGN and collectin, the mouse has more paralogs than the human. The inverse relationship is observed for the CLEC18 gene family which has three paralogs in humans and only one in the mouse. Thus, generating and using murine systems, such as full body knockout or conditional knockout mice, in future studies will provide better models to probe CLEC18 function in physiology and disease, compared to human systems in which three genetic targets have to be manipulated. Adding to the genetic complexity of the CLEC18 family, multiple paralogous sequence variants and polymorphic variants have recently been identified across the three human paralogs including a novel segmental duplication in *CLEC18A* [30]. However, despite the genetic variations in human *CLEC18* genes, the gene family is highly conserved across the Chordata phylum with the CRD segment presenting the highest degree of conservation. The high conservation in the CRD indicates that the ligands of CLEC18 are also conserved across species. The high conservation and maintenance of several CLEC18 copies in multiple species would indicate that CLEC18 is essential, however we failed to find CLEC18 orthologs in some aquatic model organisms such as zebrafish and xenopus. The reason as to why some species lack CLEC18 remains to be determined.

Compared to other canonical C-type lectins, CLEC18 have a unique CRD amino acid sequence suggesting that they bind to unique ligands. Our data now show that CLEC18 predominantely interacts with the GAG chain of proteoglycans, namely a variety of heparin, heparan sulfate, and chondroitin sulfate GAGs. So far, CLEC14A has been shown to be the only other C-type lectin that interacts with proteoglycans [23]. Whereas many studies have linked various proteoglycans and their GAGs to a wide range of kidney diseases [31, 32], further studies are needed to determine the exact role of the CLEC18-GAG axis in the context of health and disease of the kidney.

In cancer, we uncovered that CLEC18A is involved in the progression of ccRCC tumors. *CLEC18* genes are upregulated in ccRCC tumors and high CLEC18 expression correlates with improved survival of patients. When deleting *Clec18a* from murine ccRCC cells, these *Clec18a*^-/-^ mutant kidney cancer cells grew faster and presented with more neutrophil infiltration. The exact molecular mechanisms by which the secreted CLEC18 lectins and their ligands affect kidney cancer growth remain unknown. However, it is known that some of the proteoglycans and their assocaited GAGs (that we have identified as ligands of CLEC18) promote cancer. Indeed, Versican, Biglycan, Syndecans and Glypicans have all been found to increase the migratory and metastatic capabilities of different cancers through various mechanisms [33–36]. Knowing that proteoglycans can contribute to the invasiveness and metastatic spread of cancer, and that metastatic ccRCC is highly lethal [37], it is possible that CLEC18 in the tumor microenvironment reduces tumor growth and possibly metastatic spread through its interaction with various GAGs. Future studies are needed to elucidate how the CLEC18-proteoglycan axis mechanistically contributes to the growth of tumor cells, and how this pathway could be pharmacologically explored as a novel treatment for kidney cancers.

## Materials and methods

### Phylogenetic analysis

To construct a phylogenetic tree of the CLEC18 family, orthologs of the protein were collected in individual blast searches starting with human representatives in both the NCBI protein database and UniProt reference proteomes [38–40]. An E-value threshold of < 1×10^-30^was applied to identify significant matches. A selection of 37 hits covering a wide taxonomic range were then aligned using mafft (-linsi) [41], and columns with less than 20 sequences were removed (regions with long gaps). A maximum likelihood phylogenetic tree was calculated with IQ-TREE 2 v.2.2.2.6 [42], with standard model selection using ModelFinder [43]. We obtained branch supports with the ultrafast bootstrap (UFBoot2) [44] and the SH-aLRT test [45]. The tree was visualized in iTOL v6 [46] and rooted by the lamprey sequences. Branches that are supported by SH-aLRT ≥ 80% and UFboot2 ≥ 95% are indicated by a grey dot. Branch lengths represent the inferred number of amino acid substitutions per site, and branch labels are composed of gene name (if available), genus, species, and accession number. Conservation scores were calculated with Jalview using previously generated Multiple Sequence Alignments (MSA) of all found CLEC18 proteins compared to human CLEC18A. Via Jalview calculated conservation scores are based on multiple sequence alignment analysis using the AMAS method [47]. Disorder probability of the human CLEC18A protein was calculated using the IUPred3 online tool [48]. Additionally, sequences of all known human C-type lectin (CLEC) CRDs were extracted from the UniProt database, aligned using mafft (-linsi) and analyzed using Jalview. The neighbor-joining tree based on observed divergence was calculated in seaview [49] and visualized with iTOL.

The 3D structure model of CLEC18A was derived from the AlphaFold Protein Structure Database [21, 50] and visualized with ChimeraX [51]. The per residue sequence conservation score was calculated with AAcon v. 1.1. (KARLIN method, results normalized with values between 0 and 1) [52].

### RT-qPCR

Murine tissues were extracted and homogenized, and RNA isolated using an RNeasy mini kit (Qiagen). Total RNA was reverse-transcribed with the iScript cDNA Synthesis Kit (Bio-Rad). Real-time quantitative PCR was performed for murine *Clec18a* and *Gapdh* with GoTaq qPCR master mix (Promega) on the CFX384 system (Bio-Rad). Primer sequences: *Clec18a* forward: GCA GAC ACC TAC TAT GGA GCC A, *Clec18a* reverse: CAC TGT CAG TCA CCT CGT TGG T, *Gapdh* forward: GTC GGT GTG AAC GGA TTT GG, *Gapdh* reverse: GAC TCC ACG ACA TAC TCA GC.

### scRNA-sequencing re-analysis of published datasets

Processed scRNA-seq data from the the kidney cell atlas [18] was downloaded from the CZ CELLxGENE platform [53, 54] as .rds files and the Seurat objects were imported into Rstudio. All re-analysis and visualization of the kidney cell atlas was performed with Seurat v5 [55].

### CLEC18A-Fc fusion construct cloning, expression and purification

The CLEC18A-Fc fusion construct was cloned into the pCAGG_00_ccb plasmid as previously described [56] generating a construct with an inducible murine CLEC18A CRD fused to a murine IgG2a-Fc domain connected with a (GGGS)_3_ linker. Before the construct is a IL2 secretion signal allowing purification of the CLEC18A-Fc fusion from culture supernatant. The CLEC18A-fusion was expressed by transfecting the plasmid into Freestyle™ 293-F cells. Cells were seeded at 0.7×10^6^ cells/ml the day before expression and maintained after transfection at 120 rpm, 37°C, 8% CO_2_. The transfection was performed accordingly: 2 μl polyethylenimine (PEI) 25K (1 mg/ml; Polysciences) was mixed with each μg of plasmid in Opti-MEM medium (Thermo Fisher Scientific) and added to the cell suspension. 24h after transfection, the medium was topped up with EX-CELL 293 Serum-Free Medium (Sigma-Aldrich) to 20% and the culture was grown for 120h.

After expression, supernatant was harvested by centrifugating the culture at 300 x *g* for 10 min and the pellet discarded. CLEC18A-Fc fusion was purified from the supernatant with an Protein A agarose resin (Gold Biotechnology). Protein A beads in an ethanol solution were centrifuged at 150 x *g* for 5 min to remove the ethanol solution. Pure beads were washed and resuspended with 1X binding buffer (0.02 M Sodium Phosphate, 0.02% sodium azide, pH=7.0) and added into the supernatant together with concentrated binding buffer for a final concentration of 1X binding buffer. Supernatants with beads were incubated overnight 4°C on a shaking plate. After incubation, beads were collected (150 x *g*, 5 min) and washed with binding buffer. Washed beads were transferred to spin columns (G-Biosciences) and centrifuged (100 x *g*, 30 sec) to remove excess buffer. Beads were then mixed with 1 bead volume of elution buffer (100 mM Glycine-HCl, pH=2–3, 0.02% sodium azide) and incubated for 30 sec. The elution was collected into a tube containing neutralization buffer (1 M Tris, pH=9.0, 0.02% sodium azide) by centrifugation at 100 x *g* for 15 sec. The elution procedure was repeated three times, all elution fractions were pooled and the protein concentrations measured with the Pierce™ BCA ProteinAssay Kit (Thermo Fisher Scientific) using the Pierce™ Bovine Gamma Globulin Standard (Thermo Fisher Scientific).

### Flow cytometry based cell line interaction screen

All cell lines used in this study were obtained through the ATCC and cultured in DMEM (Thermo Fisher Scientific) supplemented with 10% heat inactivated FCS (Thermo Fisher Scientific), 2 mM L-glutamine (Thermo Fisher Scientific) and 1X Penicillin/Streptomyocin (Thermo Fisher Scientific). All cell lines were routinely checked for mycoplasma and validated with STR profiling.

For binding profiling, cells were harvested at approximately 80% confluency using pre-heated trypsin. Cell culture media was aspirated, cells were washed once with pre-heated PBS and then incubated with trypsin (Thermo Fisher Scientific) until detachment. Cells were harvested with pre-heated DMEM and centrifugated at 500 x *g* for 3 min. Following centrifugation, cells were re-suspended in PBS and incubated with eBioscience™ Fixable Viability Dye eFluor™ 780 (Invitrogen) for 20 min at 4°C in the dark. Cells were washed once in TSM buffer (20 mM Tris HCl, 150 mM NaCl, 2 mM CaCl_2_, 2 mM MgCl_2_) and resuspended with 10 μM CLEC18A-Fc fusion or 10 μM Fc control in TSM and incubated for 30 min at 4°C. Following primary lectin incubation, cells were washed and incubated with 1/400 F(ab’)2-Goat anti-Mouse IgG (H+L) Secondary Antibody, PE, eBioscience™ for 30 min at 4°C in the dark and then immediately analysed on a BD LSRFortessa™ Cell Analyzer flow cytometer. The data was analyzed in FlowJo (v10.4) and the binding affinity of the CLEC18A-Fc fusion lectin to various cell lines was calculated as the PE signal fold change over the control construct.

### CLEC18A-Fc fusion ligand pulldown

#### Pulldown

PymT and Ac711 cell lines were lysed through repeated freeze/thaw cycles in liquid nitrogen. After lysis, suspensions were centrifuged for 10 min at 2000 x *g* to remove cell debris and the protein concentration of the lysates were determined with the Pierce™ BCA ProteinAssay Kit (Thermo Fisher Scientific) according to the manufacturer’s instructions.

For the pulldown, 60 μl of Pierce™ Protein A/G Magnetic Beads (ThermoFisher) suspension was washed three times with PBS on a magnetic rack. After the last wash, beads were re-suspended in 500 μl PBS with 10 μg/μl CLEC18A-Fc fusion and incubated on a shaker for 2h at 4°C. In parallel, PymT and Ac711 lysates were precleared. Again, 60 μl of bead solution was washed three times with PBS and then incubated with 700 μg of lysate in 500 μl PBS. Lysates were incubated on a shaker for 2h at 4°C. After incubation, beads and lysates were washed in PBS and joined together in 500 μl TSM buffer and incubated overnight on a shaker at 4°C. After pulldown, beads were collected and washed 3 times in TSM buffer.

#### Mass spectrometry analysis of pulldown

Pulldown bead suspension were boiled with 1X Laemmli buffer (Sigma-Aldrich) and run for two minutes into a 12% NuPAGE Bis-Tris gel (Thermo Fisher Scientific) to remove impurities. Samples were subsequently gel extracted and processed further.

##### Relative peptide amount determination

Final peptide amount was determined by separation of an aliquot of each sample on an LC-UV system equipped with a monolith column. Peptide concentration was calculated based on the peak area of 100 ng of Pierce HeLa protein digestion standard (Thermo Fisher Scientific). Afterwards the peptide solution was frozen at −70°C before further processing.

##### NanoLC-MS/MS analysis

Samples were run on the nano HPLC system UltiMate 3000 RSLC nano system coupled to the Orbitrap Exploris 480 mass spectrometer, which is equipped with a NanoFlex nanospray source (Thermo Fisher Scientific). Peptides were loaded onto a trap column (Thermo Fisher Scientific, PepMap C18, 5 mm × 300 μm ID, 5 μm particles, 100 Å pore size) at a flow rate of 25 μl/min using 0.1% TFA as mobile phase. After 10 min, the trap column was connected in series with the analytical column (Thermo Fisher Scientific, PepMap C18, 500 mm × 75 μm ID, 2 μm, 100 Å). The analytical column further was connected to PepSep sprayer 1 (Bruker) equipped with a 10 μm ID fused silica electrospray emitter with an integrated liquid junction (Bruker, PN 1893527). Electrospray voltage was set to 2.4 kV. Peptides were eluted using a flow rate of 230 nL/min and a 120 min gradient. The gradient starts with mobile phases A and B: 98% A (water/formic acid, 99.9/0.1, v/v) and 2% B (water/acetonitrile/formic acid, 19.92/80/0.08, v/v/v), increases to 35% B over the next 120 min. This is followed by a gradient increase to 90% B within the next 5 min, stays there for 5 min and decreases again within 2 min back to a gradient of 98% for A and 2% for B to equilibrate at 30°C. The Orbitrap Q-Exactive HF-X mass spectrometer was operated by a mixed MS method which consisted of one full scan (m/z range 380-1 500; 15 000 resolution; AGC target value 3 x 10^6^). Maximum injection time was set to 800 ms.

##### Data analysis

For peptide identification, RAW-files were loaded into Proteome Discoverer (version 2.5.0.400, Thermo Scientific). All created MS/MS spectra were searched using MSAmanda v2.0.0.19924 (Dorfer et al., 2014). As a first step RAW-files were searched against the uniprot_reference_mouse_2023-03-06.fasta (21 928 sequences; 11 722 545 residues), PD_Contaminants_TAGs_v20_tagsremoved.fasta, tags_v11.fasta data bases using following search parameters: The peptide mass tolerance was set to ±10 ppm and the fragment mass tolerance to ±10 ppm. The maximal number of missed cleavages was set to 2. The result was filtered to 1% FDR on protein level using the Percolator algorithm integrated in Thermo Proteome Discoverer. Additionally a sub-database was generated for further processing. As a second step RAW-files were searched against the created sub-database. The following search parameters were used: Iodoacetamide derivative on cysteine was set as a fixed modification, oxidation on methionine, phosphorylation on serine, threonine and tyrosine, deamidation on asparagine and glutamine, pyro-glu from q on peptide N-terminal glutamine, acetylation on protein N-Terminus were set as variable modifications. Monoisotopic masses were searched within unrestricted protein masses for tryptic enzymatic specificity. The peptide mass tolerance again was set to ±10 ppm and the fragment mass tolerance to ±10 ppm. Maximal number of missed cleavages was set to 2. The result was filtered to 1% FDR on protein level using Percolator algorithm integrated in Thermo Proteome Discoverer. Additional high quality filtering by setting a minimum MS Amanda Score of 150 on PSM level was applied.

Protein areas were quantified using IMP-apQuant [57] by summing unique and razor peptides and applying intensity-based absolute quantification (iBAQ) [58] with subsequent normalisation based on the MaxLFQ algorithm [59]. Proteins were filtered to be identified by a minimum of 2 PSMs in at least one sample and identified proteins were pre-filtered to contain at least three quantified peptide groups. Pulldown hits are shown in **Supplementary Table 1**.

#### AlphaFold2 Multimer interaction screen

AlphaFold2 Multimer [20–22] was impleneted and used to predict interactions between CLEC18A, CLEC18B and CLEC18C and putative interaction partners. A list of 379 kidney and clear cell renal cell carcinoma specific proteins was generated by: extracting (i) proteins that in the literature are defined as proximal tubule specific (expression site of CLEC18A) [18, 60, 61], (ii) proteins that in the human protein atlas [62] are defined as kidney enriched and/or only found in kidney, and (iii) proteins that were angiotensin related [63], were pathway signature associated [64], had high mutational frequency [65], were associated with a ccRCC inflammatory state [66], had high recurrent repeat expansions in ccRCC [67], or proteins that in the cBioPortal [68] were mutated in more than 5% of all ccRCC cases and/or correlated (Spearman’s correlation > 0.5) with *CLEC18A* expression. After assembling the list of targets redundancies were removed. CLEC18A, CLEC18B and CLEC18C were all screened against the list of proteins resulting in 1110 pairwise PPI predictions using a custom script on a local CPU and GPU cluster using MMseqs (https://github.com/soedinglab/MMseqs2) for local Multiple Sequence Alignment creation and colabfold (https://github.com/sokrypton/ColabFold) for structure prediction with 5 models per prediction and omitting structure relaxation. Hits from the CLEC18A-Fc fusion pulldown experiment were analyzed in the same way.

#### GlycoArray

Glycan arrays were printed as previously described by [69]. In brief, 254 glycans (**Supplementary Table 2**) were printed using an Arrayjet Mercury Microarray printer at a concentration of 1000 µM onto sciCHIP epoxy slides (Scienion). Slides were neutralised, rinsed in 100% ethanol and stored at 4°C under vacuum until use. For all array work, proteins are diluted in array PBS (PBS, 1.8 mM MgCl_2_, 1.8 mM CaCl_2_, pH 7.4), filtered through 0.22 µm filter and degassed prior to use. 2 µg of Fc control and CLEC18A-Fc fusion were incubated with AlexaFluor647 rabbit anti-mouse and goat anti-rabbit IgG (Invitrogen) at a molar ratio of 4:2:1 in array PBS and preincubated on ice for 10 mins at a total volume of 65 µl. A 65 µl geneframe was placed onto the slide surface and the incubation mix was subsequently added and sealed with a coverslip. The sample was incubated for 15 mins at room temperature in the dark. After the incubation time, the slide was immersed in array PBS and the geneframe and coverslip were removed. The slides were washed briefly for two mins with gentle shaking. The slides were dried by centrifugation at 200 x *g* for 4 min. The slide was then scanned using the 647 nm laser and 677/45 filter, with low laser power and 50% PMT gain settings. The acquired image was then analysed using the Mapix software, overlaying the image with the map (Gal) file. The experiment was repeated three times, and binding was classified as positive when the average RFU (relative fluorescence units) of a specific structure had a value above mean background (defined as the average background fluorescence plus 3 standard deviations) and had a P value of < 0.005 (student’s t-test). A pre-experimental image was acquired to rule out any autofluorescence of the printed glycans.

#### TCGA kidney cancer data analysis

TCGA RNA sequencing data from tumor and normal tissue for TCGA-KIRC, TCGA-KIRP and TCGA-KICH was downloaded through the GDC portal. Differential expression analysis between tumor and matched normal tissue was performed with DESeq2 (RRID:SCR_015687, v1.22.2). Significantly upregulated or downregulated genes were defined as genes with ≥10 counts, and Benjamini-Hochberg adjusted Wald p values of p < 0.05.

### *Clec18a* knockout generation

gRNAs for murine *Clec18a* were designed using CHOPCHOP [70], synthesized with appropriate overhangs and cloned into the pSpCas9(BB)-2A-Puro (PX459) V2.0 plasmid [71]. Cells were transfected using the MaxCyte® STX electroporation system. For this, 10^6^ cells per cell line were prepared by washing with PBS and HyClone Electroporation Buffer (MaxCyte® once. Then, RAG cells were resuspended with 100 µl HyClone Electroporation Buffer (MaxCyte) and 20 µL of PX459 plasmid containing the *Clec18a* gRNA. The mix containing cells and plasmid was transferred into an OC-100 cuvette and electroporated using the “Renca” program from MaxCyte. Electroporated cells were then transferred into a 10 cm dish and incubated for 30 min at 37°C. Then, they were resuspended with 10 ml of supplemented DMEM media and incubated over night at 37°C. The next day for selecting transfected cells 2 µg/ml of puromycin was added for 24 hours. After seeded cells reached confluency single cell clones were seeded in two 96-well plates per cell line using the FACS Aria III cell sorter. Individual clones were a region flanking the mutation site amplified using PCR and the PCR fragment was purified and subjected to Sanger sequencing (Vienna BioCenter inhouse service). The editing efficiency was assessed using TIDE analysis [72]. Clones with a predicted efficiency of >95% and with a frameshift mutation were expanded and validated using RT-qPCR.

### *Clec18a* overexpression generation

E0771 cells overexpressing *Clec18a* were generated using a lentiparticle construct pre-assembled by OriGene. E0771 cells were transduced with lentiparticles containing containing *Clec18a* overexpression plasmid or an empty control plasmid, both with puromycin resistance. According to the manufacturer’s instructions, on the first day 0.24 x 10^6^ cells were seeded in each well of a 6-well plate, whereas two wells were seeded per cell line, one for each condition and incubated for 20 hours at 37°C. The next day for viral infection of the cells lentiviral particles were added according to a multiplicity of infection (MOI) of 10, cell culture media and 8 µg/mL of polybrene. Cells were then incubated for 18 hours at 37°C. Afterwards lentivirus containing medium was removed and exchanged with normal complete DMEM. After cell line generation, E0771 cells were maintained in media containing 1 µg/ml puromycin.

#### Mouse studies

##### Mus musculus

All mice used in this study were bred and maintained at the Comparative Medicine Mousehouse at the Vienna BioCenter. The mice were maintained in pathogen free environments with regular health screenings in a 12 hour light-dark cycle. Food and water was provided *ab libitum*. All mouse experiments were approved by the Bundesministerium für Wissenschaft, Forschung und Wirtschaft (BMWFW, project 2023-0.517.898), and carried out according to EU-directive 2010/63/EU.

#### Subcutaneous tumor injection

Clear cell renal cell carcinoma cells (RAG) were injected subcutaneously into the left flank and breact cancer cells (E0771) were injected orthotopically into the right inguinoabdominal mammary fat pad under anaesthesia with xylasol and ketasol. Per mouse, 500000 RAG cells or 250000 E0771 cells were injected. RAG cells did not grow properly in wildtype syngenic BALB/c mice so all mouse experiments were performed in *Rag2*^-/-^ *Il2rg*^-/-^ immunodeficient mice (The Jackson Laboratory). Tumor size was measured every 3-4 days and the volume was calculated as 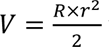 where R is the longest diameter and r is short diamater.

#### Immunohistochemistry

Tumors were harvested as mice reached human endpoint and fixed in neutral buffered formalin (Sigma-Aldrich) for 24-48h, room temperature. Following fixation, tumors were washed in PBS and embedded in paraffin and sectioned at 2 μm thickness. Sections were mounted on glass slides (Permaflex plus adhesive slides, Leica). Slides were deparaffinized and rehydrated with a Gemini AS automatic stainer (Fisher Scientific). Rehydrated slides were subjected to heat-induced epitope retrieval with pH 6 sodium citrate buffer, followed by endogenous peroxidase inactivation with 3% H_2_O_2_ (Sigma-Aldrich) and subsequently blocked in 5% BSA in TBS-T with TBS washes inbetween each step. Slides were incubated with primary rabbit anti-Ly6G or rabbit anti-F4/80 antibodies (Cell Signaling Technology) for 2h at room temperature and subsequently treated with the a rabbit detection system (DCS) according to the manufacturers instructions. Chromogenic reaction was induced with a DAB substrate kit (Abcam) and hematoxylin counter taining was performed with the Gemini AS automatic stainer. Slides were mounted with cover slips by Tissue-TEK GLC (Sakura Finetek) and scanned with Slide Scanner Pannoramic 250 (3DHistech Ltd).

### RNAScope *in situ* hybridization

Brains were extracted from 10 week old wildtype C57B6J mice and fixed overnight in 4% paraformaldehyde followed by dehydration and paraffin embedding. Following embedding, 3.5 μm sections were cut throughout the brain and *in situ* hybridized with a custom designed RNAScope probe for murine *Clec18a* by ACD biosciences. *In situ* hybridization was performed according to manufacturers instructions. The sections were pretreated for 15 minutes with Target Retrieval Solution and for 30 minutes with Protease Plus. After hybridization, the mRNA signal was amplified and visualized with an alkaline phosphatase– based red substrate.

### Statistical analysis

All statistical analysis were performed in Prism 10 (GraphPad) or in RStudio (Posit). When comparing large omics datasets p values were calculated with limma-moderated Benjamini– Hochberg-corrected two-sided t-test after data processing for proteomics, and Benjamini-Hochberg adjusted Wald p values for TCGA data. For comparisons of individual markers between groups, the distribution of the data was initially determined by Shapiro-Wilk normality test. Statistical testing was then performed with Mann-Whitney U-tests (non-normal data) or two-sided t-tests (normal data). Tumor time course significance was assessed with two-way ANOVA with Šídák’s multiple comparisons test.

## Disclosure statement

The authors have no conflicts of interest to declare.

## Author contribution

G.J. and J.M.P conceived the study. G.J. performed and designed experiments with input and help from all co-authors as follows: M.H. with cell line generation, mouse experiments and AlphaFold2 Multimer screens, L.H-T with glycoarray experiments, S.M. and T.O. with pulldowns and mass spectrometry, I.S. with mouse experiments, D.H. with recombinant CLEC18A-Fc generation, M.N. with TCGA analysis and A.S. with phylogeny analysis.

## Supporting information

Supplementary table 1 Lectin pulldown raw data

Supplementary table 2 Glycoarray raw data

## Acknowledgements

We thank all members of our laboratories for their support and help. The RNA-FISH and immunohistochemistry was performed by the Histology Facility at Vienna BioCenter Core Facilities (VBCF), member of the Vienna BioCenter (VBC), Austria. Proteomics analyses were performed by the Proteomics Facility at IMP/IMBA/GMI using the VBCF instrument pool. The YUMM cell lines were a gift from the Obenauf lab. G.J. is supported by a DOC fellowship from the Austrian Academy of Sciences. S.M. received funding from the European Union’s Horizon 2020 research and innovation programme under the Marie Sklodowska-Curie grant agreement No 841319 and the ESPRIT-Programme of the Austrian Science Fund (FWF, Project number: ESP 166). The lab of J.M.P. received funding from the Medical University of Vienna, the Austrian Academy of Sciences, the T. von Zastrow foundation, the Fundacio La Marato de TV3 (202125-31), and the Canada 150 Research Chairs Program F18-01336. We also gratefully acknowledge funding by the German Federal Ministry of Education and Research (BMBF) under the project “Microbial Stargazing - Erforschung von Resilienzmechanismen von Mikroben und Menschen” (Ref. 01KX2324).

## Data availability

Data for the glycoarray and hits from the pulldown experiment are available as supplementary tables. All other raw data is available from the lead authors upon request.

## Conflict of interest

The authors have no conflict of interest to declare

## Figures and Figure legends

**Supplementary Figure 1.**
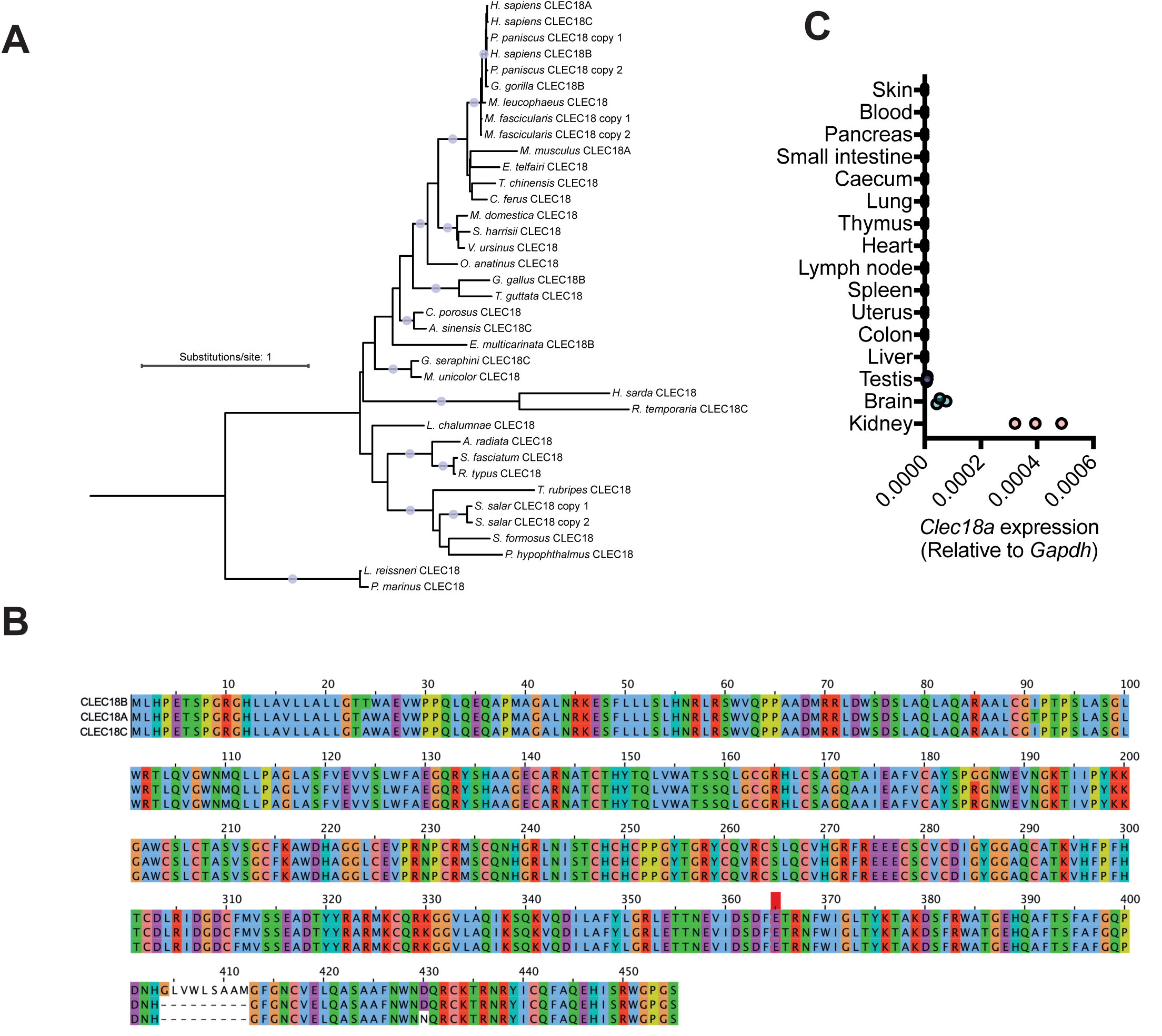
Conservation and expression of CLEC18A. **A.** Maximum Likelihood phylogenetic tree based on a multiple sequence alignment of selected CLEC18 protein sequences. Branches that are supported by SH-aLRT ≥ 80% and UFboot2 ≥ 95% are indicated by a grey dot. **B.** Multiple sequence alignment of human CLEC18A, CLEC18B and CLEC18C proteins colored according to the Clustal scheme. **C.** *Clec18a* expression in mouse organs determined by RT-qPCR. Organs were always harvested from there individual mice.

**Supplementary Figure 2.**
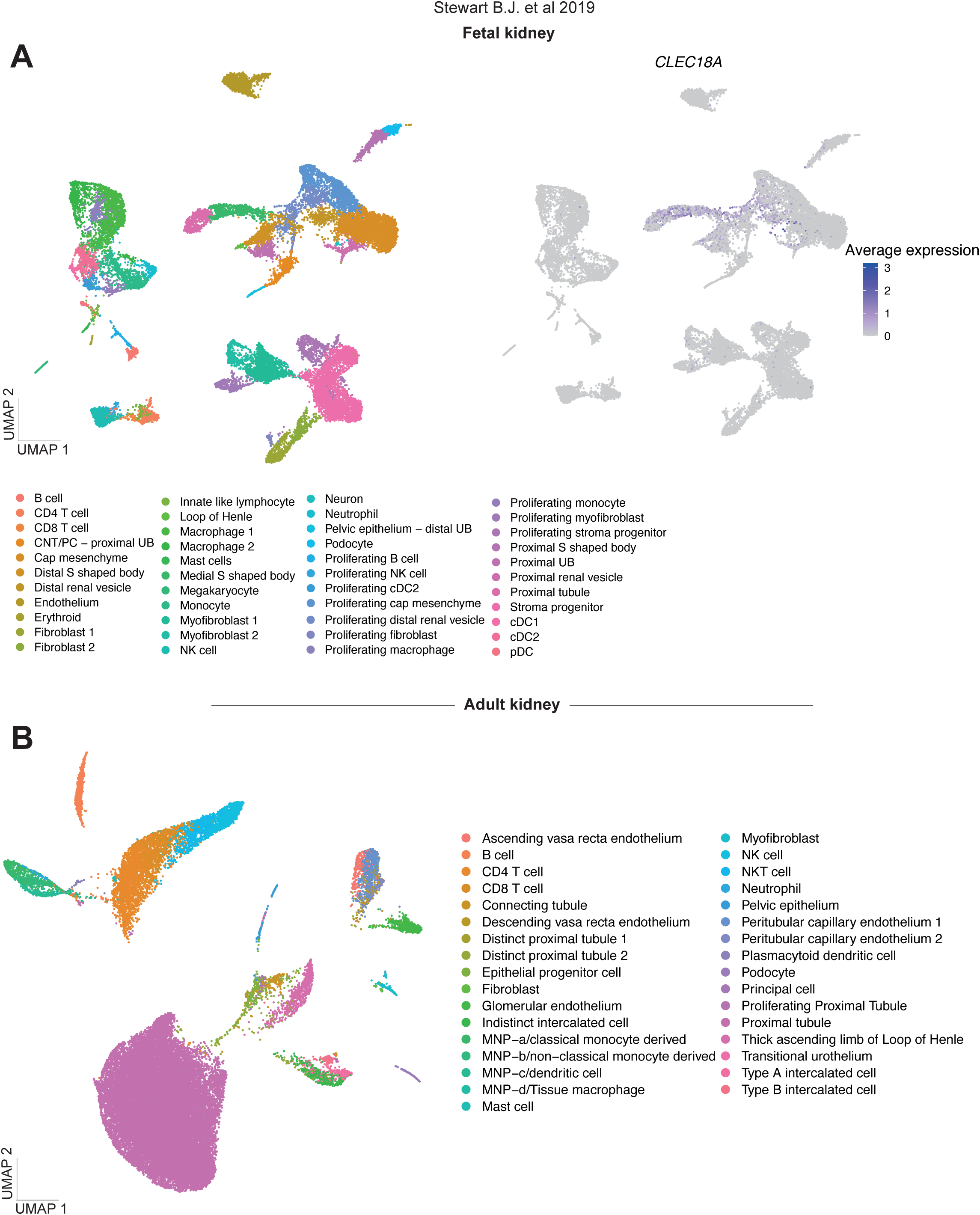
scRNA-seq clustering of fetal kidney and adult kidney cells as accoridng to Stewart B.J. et al 2019 [18]. **A.** Re-analysis of scRNA-seq data from the kidney cell atlas [18] showing clustering of fetal kidney cells and *CLEC18A* expression in the full fetal kidney. Abbreviations: UMAP = Unifold manifold approximation and projection, CNT = Connecting tubule, PC = Principal cell, UB = Ureteric bud, NK cell = Natural killer cell, cDC1 = Conventional dendritic cell type 1, cDC2 = Conventional dendritic cell type 2, pDC = Plasmacytoid dendritic cell. **B.** Re-analysis of scRNA-seq data from the kidney cell atlas [18] showing clustering of adult kidney cells. Abbreviations: UMAP = Unifold manifold approximation and projection, MNP = Mononuclear phagocyte, NK cell = Natural killer cell.

**Supplementary Figure 3.**
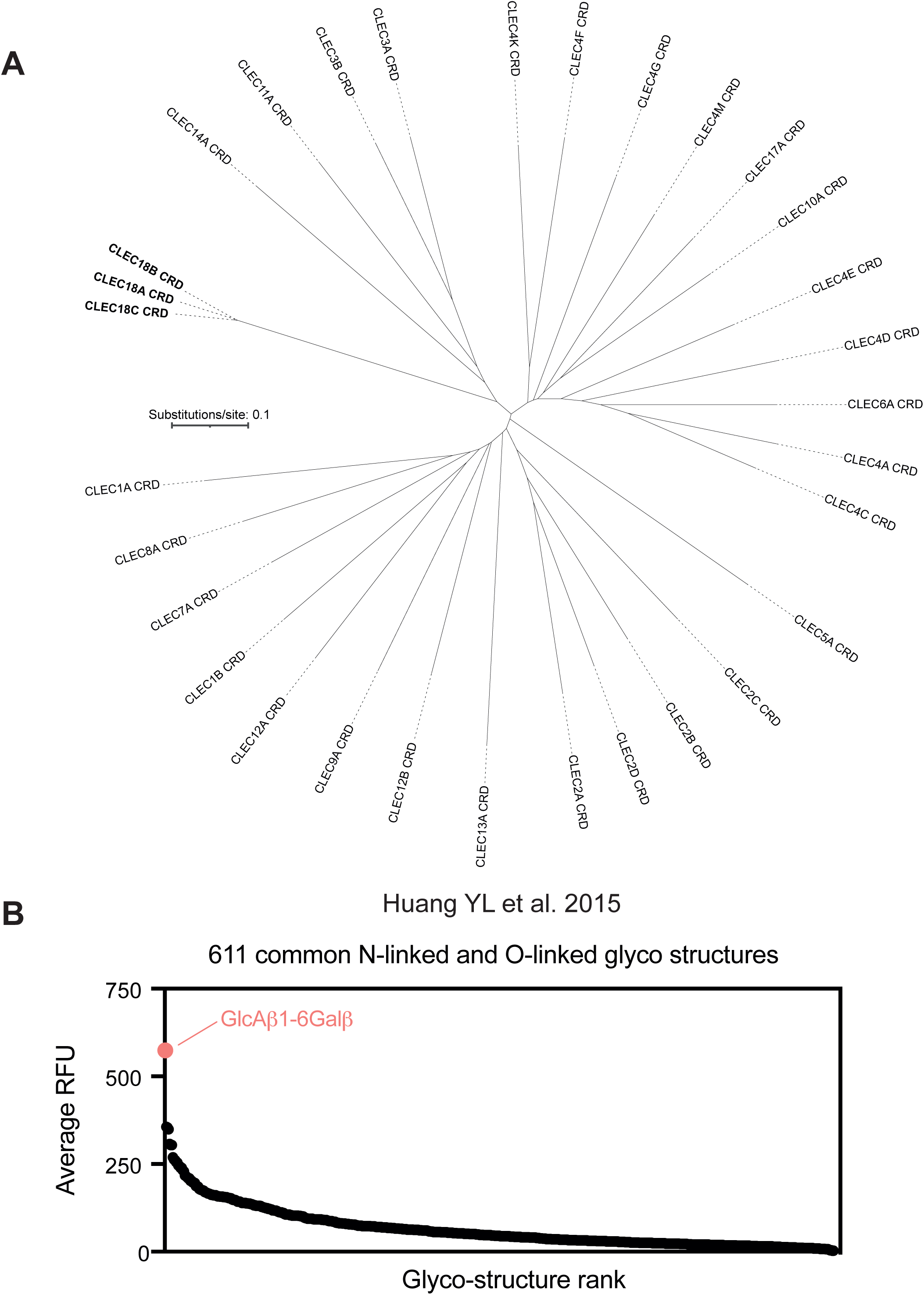
CLEC18 CRDs have unique phylogeny and ligands. **A.** Unrooted neighbourhood joining phylogenetic tree based on the multiple sequence alignment of all included C-type lectin CRD sequences. Abbreviations: CRD = Carbohydrate recognition domain. **B.** Re-analysis of a recombinant CLEC18A used on a GlycoArray containing 611 common N-linked and O-linked glyco structures [13].

**Supplementary Figure 4.**
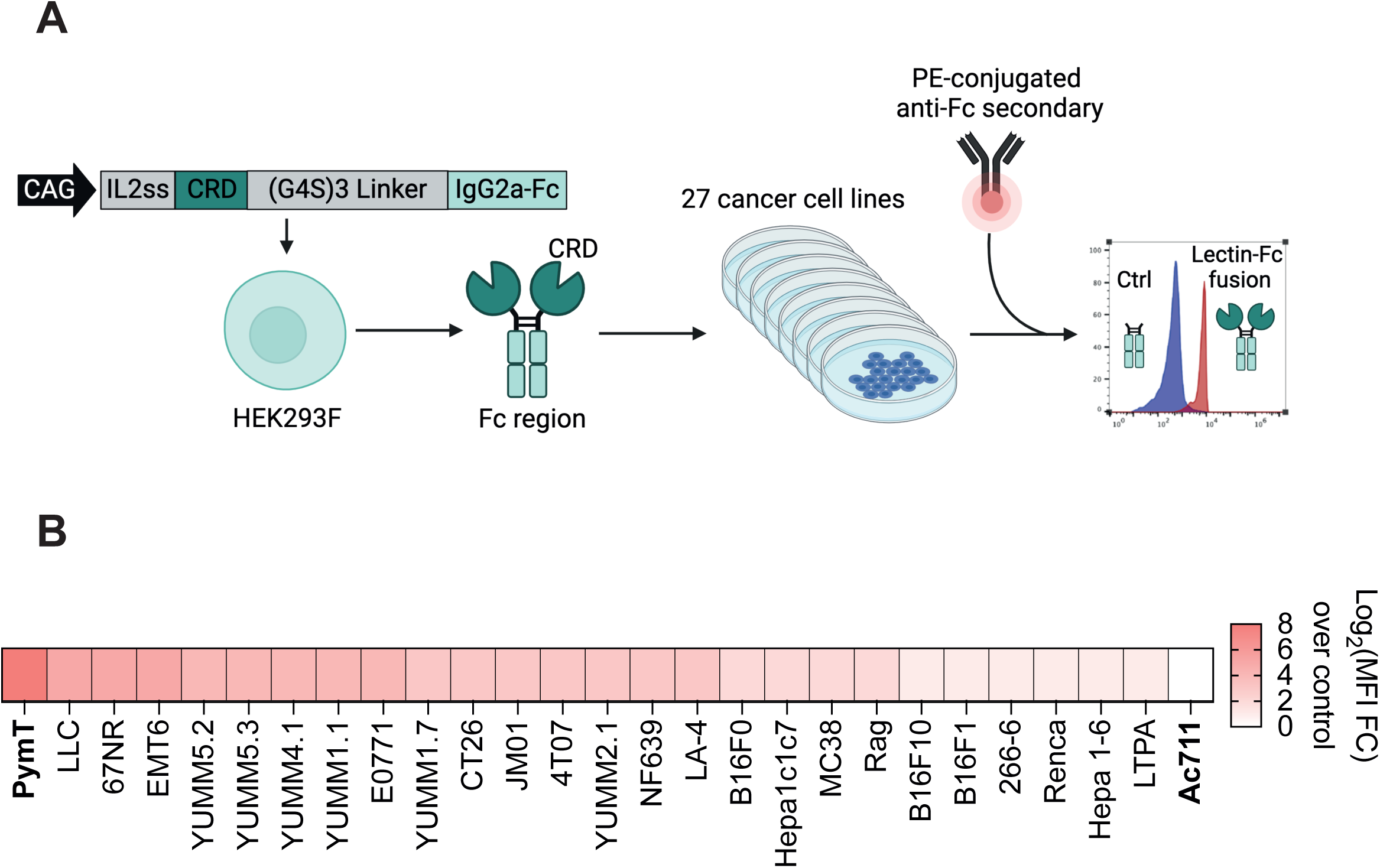
CLEC18A-Fc binds cell lines with varying efficiency. **A.** Schematic of vector used for CLEC18A-Fc expression, subsequent expression and purification, and flow cytometry based readout used to identify cell lines CLEC18A-Fc binds to. Schematic created with BioRender.com. **B.** Binding affinity of CLEC18A-Fc against a selection of cancer cell lines.

**Supplementary Figure 5.**
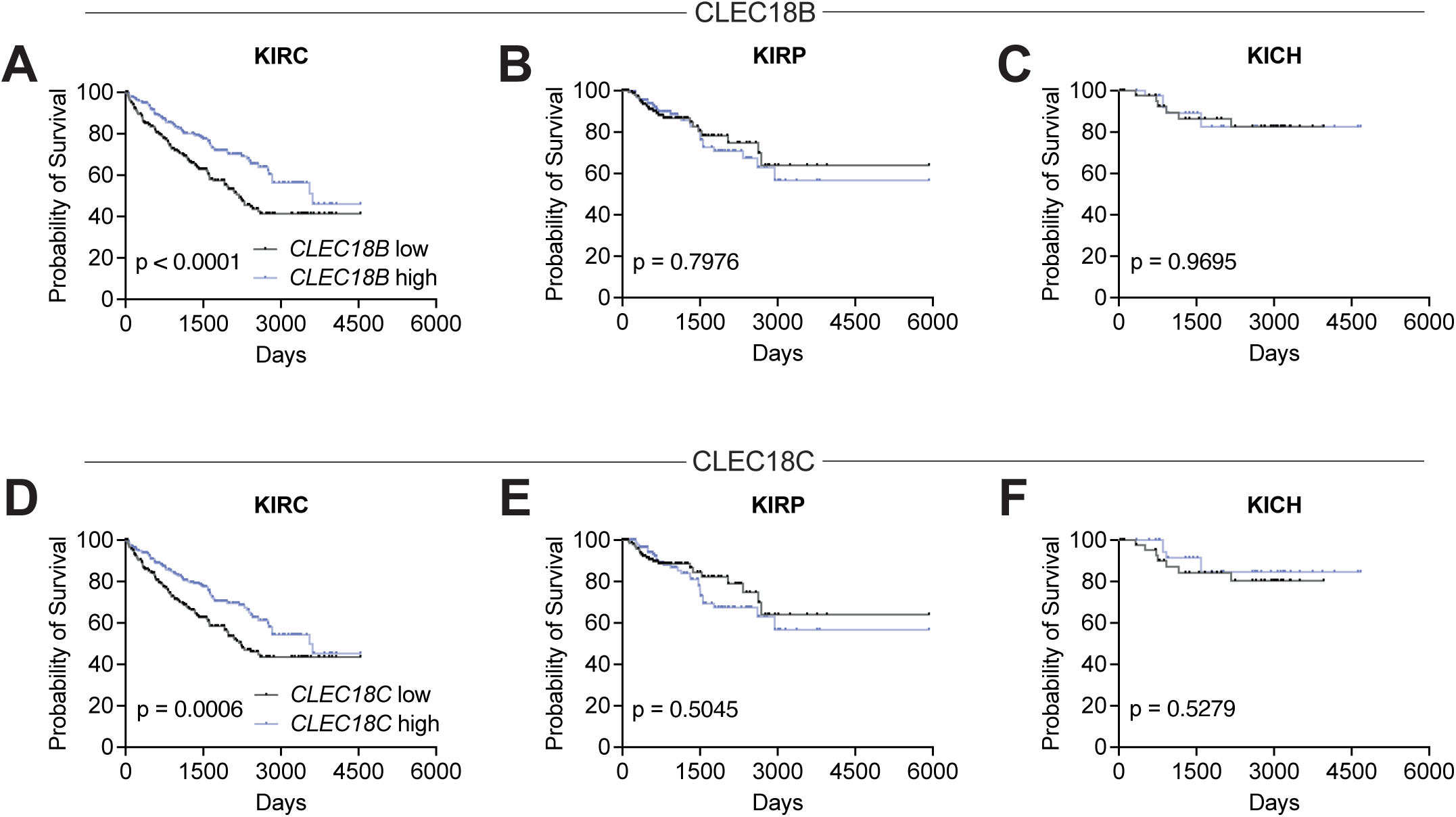
CLEC18 exclusively promotes survival in ccRCC/KIRC. **A.** Kaplan meier survival curve for high vs low *CLEC18B* expression (high and low expression attributed based on the median) in KIRC. P value calculated with a Log rank test. Abbreviations: KIRC = Clear cell renal cell carcinoma. **B.** Kaplan meier survival curve for high vs low *CLEC18B* expression (high and low expression attributed based on the median) in KIRP. P value calculated with a Log rank test. Abbreviations: KIRP = Papillary renal cell carcinoma. **C.** Kaplan meier survival curve for high vs low *CLEC18B* expression (high and low expression attributed based on the median) in KICH. P value calculated with a Log rank test. Abbreviations: KICH = Chromophobe renal cell carcinoma. **D.** Kaplan meier survival curve for high vs low *CLEC18C* expression (high and low expression attributed based on the median) in KIRC. P value calculated with a Log rank test. Abbreviations: KIRC = Clear cell renal cell carcinoma. **E.** Kaplan meier survival curve for high vs low *CLEC18C* expression (high and low expression attributed based on the median) in KIRP. P value calculated with a Log rank test. Abbreviations: KIRP = Papillary renal cell carcinoma. **F.** Kaplan meier survival curve for high vs low *CLEC18C* expression (high and low expression attributed based on the median) in KICH. P value calculated with a Log rank test. Abbreviations: KICH = Chromophobe renal cell carcinoma.

## Notes

### Competing Interest Statement

The authors have declared no competing interest.

### Summary of Updates

Updated abstract (correction of typos, sulfonated has been corrected to sulfated) Added keywords

